# Intranasal type I interferon treatment is beneficial only when administered before clinical signs onset in the SARS-CoV-2 hamster model

**DOI:** 10.1101/2021.02.09.430458

**Authors:** Pierre Bessière, Marine Wasniewski, Evelyne Picard-Meyer, Alexandre Servat, Thomas Figueroa, Charlotte Foret-Lucas, Amelia Coggon, Sandrine Lesellier, Frank Boué, Nathan Cebron, Blandine Gausserès, Catherine Trumel, Gilles Foucras, Francisco J Salguero, Elodie Monchatre-Leroy, Romain Volmer

## Abstract

Impaired type I interferons (IFNs) production or signaling have been associated with severe COVID-19, further promoting the evaluation of recombinant type I IFNs as therapeutics against SARS-CoV-2 infection. In the Syrian hamster model, we show that intranasal administration of IFN-α starting one day pre-infection or one day post-infection limited weight loss and decreased viral lung titers. By contrast, intranasal administration of IFN-α starting at the onset of symptoms three days post-infection had no impact on the clinical course of SARS-CoV-2 infection. Our results provide evidence that early type I IFN treatments are beneficial, while late interventions are ineffective, although not associated with signs of enhanced disease.

**One Sentence Summary:** The timing of type I interferon treatment is a critical determinant of its efficacy against SARS-CoV-2 infection.

## Main Text

Type I interferons (IFNs) are major antiviral cytokines and their finely tuned production is critical for host protection against viruses (*1*). *In vitro* studies demonstrated that SARS-CoV-2 was very sensitive to the antiviral effects of type I IFN (*2–4*). In addition, development of severe COVID-19 was shown to correlate with decreased type I IFNs production or impaired type I IFN signaling (*5–9*). In agreement with these observations, recombinant type I IFNs are being tested in a number of clinical trials to treat COVID-19 patients (*10–12*). However, the design and interpretation of these clinical trials need to consider that the timing of type I IFN treatment may be critical for its efficacy and safety against SARS-CoV-2 (*13, 14*). Indeed, studies in SARS-CoV and MERS-CoV infected mice demonstrated that type I IFN-treatment was beneficial when administered early, while it was deleterious when administered at later stages of infection (*15, 16*). How the timing of type I IFN treatments modulate their clinical efficacy against SARS-CoV-2 is currently unknown and needs to be tested in an animal model.

To address this question, we designed a study that evaluated the prophylactic and therapeutic efficacy of intranasally administered recombinant universal IFN-α (rIFN) against SARS-CoV-2 infection in Syrian hamsters (Fig. 1A). Hamsters intranasally infected with a high dose SARS-CoV-2 develop clinical disease caused by lung pathology, which closely mirrors severe human COVID-19 (*17, 18*). In a preliminary experiment, we observed a significant upregulation of the expression of the type I IFN stimulated gene (ISG) Mx2 in the nasal turbinates, lung and spleen of hamsters treated intranasally with rIFN, demonstrating that this molecule was active in hamsters (Fig. S1). Following challenge with 10^4^ TCID_50_ of SARS-CoV-2, no protection from weight loss was observed in the IFN-late group, for which treatment was initiated at the onset of clinical signs, when infected animals started to significantly lose weight three days post-infection (Fig. 1B). By contrast, we observed a significant protection from weight loss in the IFN-pre group (prophylactic treatment initiated 16 hours before infection) and in the IFN-early group (treatment initiated at one day post-infection) compared to the placebo group (Fig. 1B). The protection from weight loss in the IFN-pre and in the IFN-early groups was not associated with a reduction of viral excretion level or duration, as viral RNA levels measured by RT-qPCR from oropharyngeal swabs were similar in all groups (Fig. 1C). In agreement with this observation, subgenomic viral RNA levels in the nasal turbinates were similar in all groups (Fig. S2). As SARS-CoV-2 respiratory disease is due to lower respiratory tract damage, we analyzed viral load in the lungs. We detected a reduction of pulmonary viral subgenomic RNA levels and infectious viral titers in all the IFN-treated groups at day 5 post-infection, compared to the placebo group, which reached statistical significance in the IFN-early group only (Fig. 1 D&E).

**Fig. 1.**
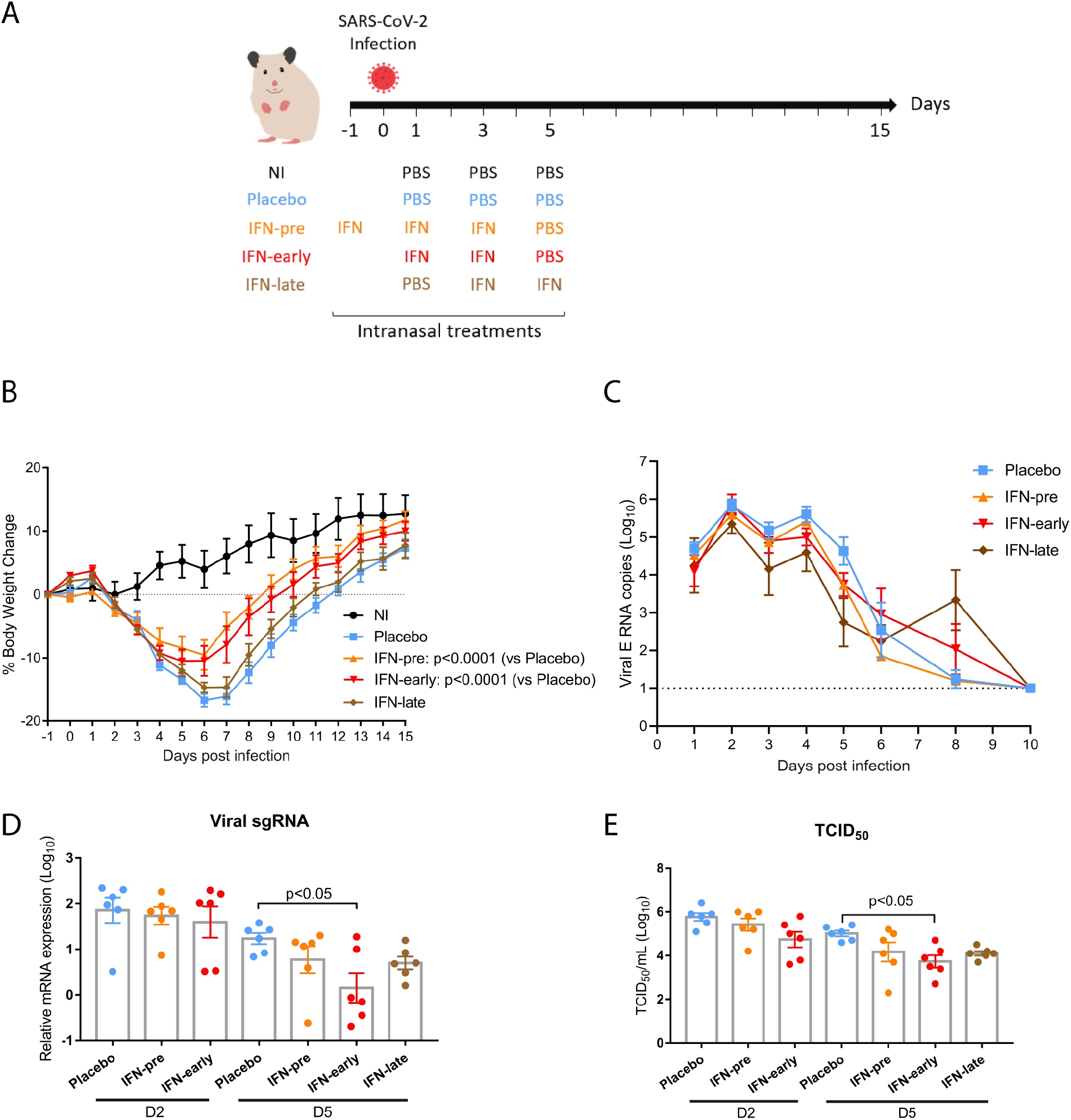
Impact of type I IFN-α treatment on SARS-CoV-2-induced weight loss and viral titers. **(A)** Overview study design. **(B)** Percentage of body weight change with weight measured day 1 pre-infection before the IFN-pre treatment set as the reference weight (6 animals per group). **(C)**. Viral genomic RNA in oropharyngeal swabs (6 animals per group). The dotted line indicates limit of detection. **(D** and **E)** Lungs viral titers determined by RT-qPCR targeting viral sgRNA relative to the housekeeping genes RPL18 and RPS6KB1 **(D)** and by TCID_50_ **(E)**. D2: day 2 post infection; D5: day 5 post infection. Results are expressed as means ± SEM.

Evaluation of the respiratory tract from infected animals revealed a mild to moderate bronchointerstitial pneumonia at day 2 post-infection, progressing to moderate/severe with lung consolidation at day 5 and resolving at day 15 with only small lesioned areas remaining, as previously observed (*18*). The lesions were characterized by infiltrates of macrophages and neutrophils, with fewer lymphocytes and plasma cells (Fig. 2A & Fig. S3). A reduction of the lung pathology scores was observed in the IFN-treated groups compared to the placebo group (Fig. 2B). RNAScope *in situ* hybridization (ISH) was used to determine the localization of viral RNA in the lungs of infected animals. Viral RNA was observed in bronchial and bronchiolar epithelial cells and in regions of inflammatory infiltrates at day 2 post-infection (Fig. S3). The viral RNA positive area diminished at day 5 and coincided with inflammatory infiltrates. Quantification of viral RNA positive area revealed a slight non-statistically significant reduction of viral RNA in the IFN-pre and in the IFN-early groups at day 2 and 5 post-infection compared to the placebo group (Fig. 2C). Mx1 is an ISG and its level of expression is a good indicator of the levels of type I and type III IFN produced locally (*19*). Mx1 protein was upregulated in the lungs of infected hamsters, as detected by immunohistochemistry, and the percentage of Mx1 positive lung was equivalent in placebo and IFN-treated hamsters (Fig. 2D & Fig. S3). Finally, hematological analyses revealed a modest lymphocytopenia in SARS-CoV-2 infected hamsters, with no difference between the IFN-treated groups and the placebo group (Fig. S4).

**Fig. 2.**
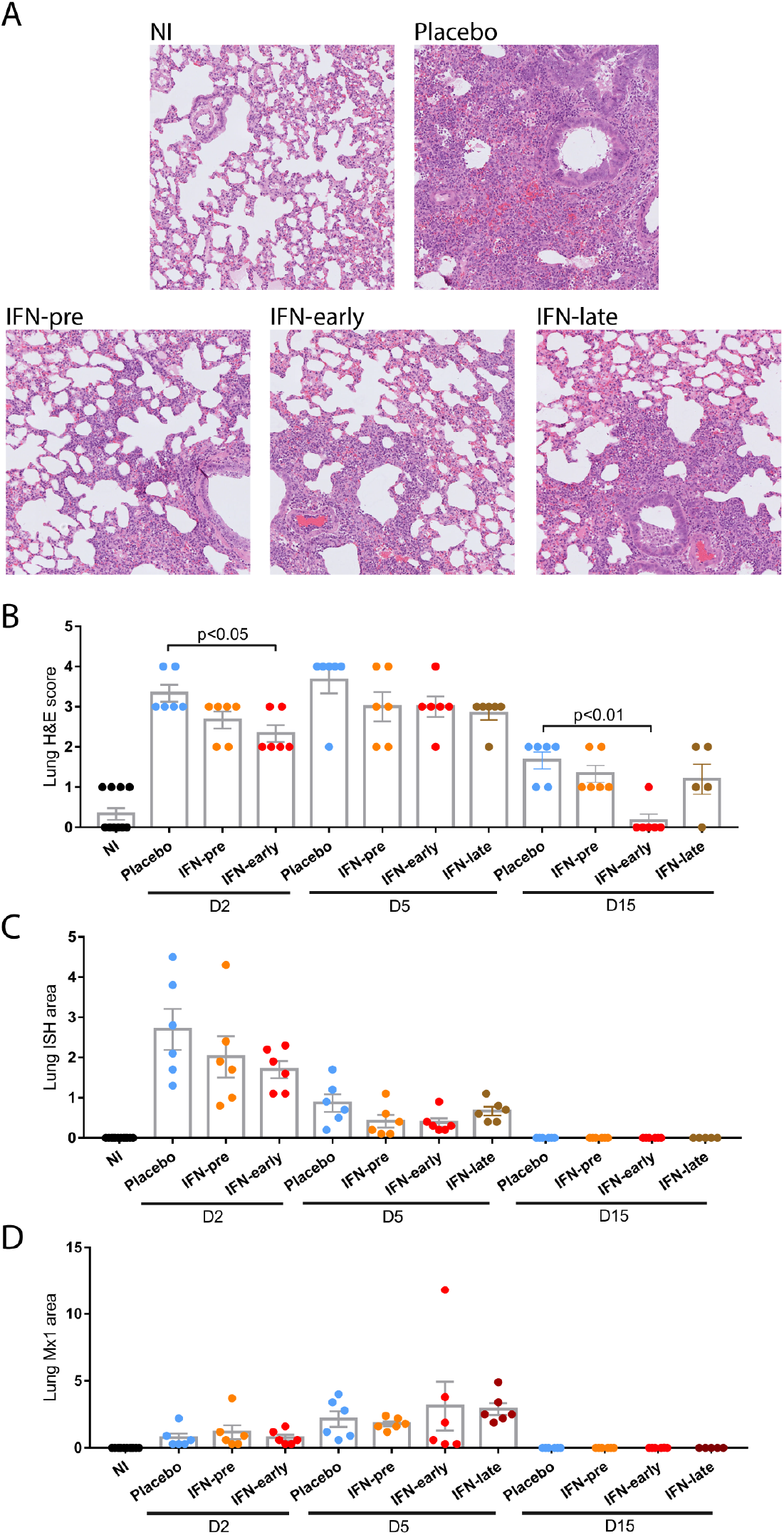
Histopathological analysis of the impact of type I IFN-α treatment. **(A)** Representative pictures were selected to display the pathology from haematoxylin and eosin (H&E) stained lung section from animals at day 5 post infection. **(B)** Severity of lung pathology based on lesional scores evaluated from haematoxylin and eosin (H&E) stained lung section. **(C)** Quantification of percent lung area positive for viral RNA in lung sections stained with RNAScope *in situ* hybridization (ISH). **(D)** Quantification of percent lung area positive for Mx1 protein detected by immunohistochemistry (IHC). D2: day 2 post infection; D5: day 5 post infection; D15: day 15 post infection. Results are expressed as means ± SEM.

To explore the consequences of type I IFN administration on the immune response to SARS-CoV-2 infection, we analyzed the expression of immune markers gene expression from the lungs of animals euthanized at day 2 and 5 post-infection. Compared to the non-infected animals, all the infected groups presented a significant upregulation of the ISGs Mx2 and ISG15 and of the C–X–C motif chemokine ligand 10 (CXCL10) messenger RNA (mRNA) expression at day 2 and 5 post-infection, with no difference between the placebo and the IFN-treated groups (Fig. 3A). The mRNA expression levels of IFN-γ, and the interleukins (ILs) IL-10 and IL-6 were also significantly upregulated in the infected animals at day 5 post-infection. Similar results were obtained for other immune markers analyzed by RT-qPCR in the lung (Fig. S5), nasal turbinates (Fig. S6) and spleen (Fig. S7). We also measured the protein levels of chemokine and cytokines either in the lungs or plasma using a commercial enzyme-linked immunosorbent assay (ELISA) directed against hamster IL-6 or a custom-developed hamster multiplex assay. Compared to noninfected animals, we detected an upregulation of CXCL10 and IL-10 protein levels in the lung of all infected groups, with no difference between the placebo and the IFN-treated groups (Fig. 3B). We detected a reduction of lung IL-1β levels in IFN-treated groups compared to placebo. Interestingly, lung IL-6 protein level and plasmatic chemokine ligand 2 (CCL2) and tumor necrosis factor-α (TNF-α) protein levels were upregulated in the IFN-late group, compared to the IFN-pre and IFN-early group (Fig. 3B&C).

**Fig. 3.**
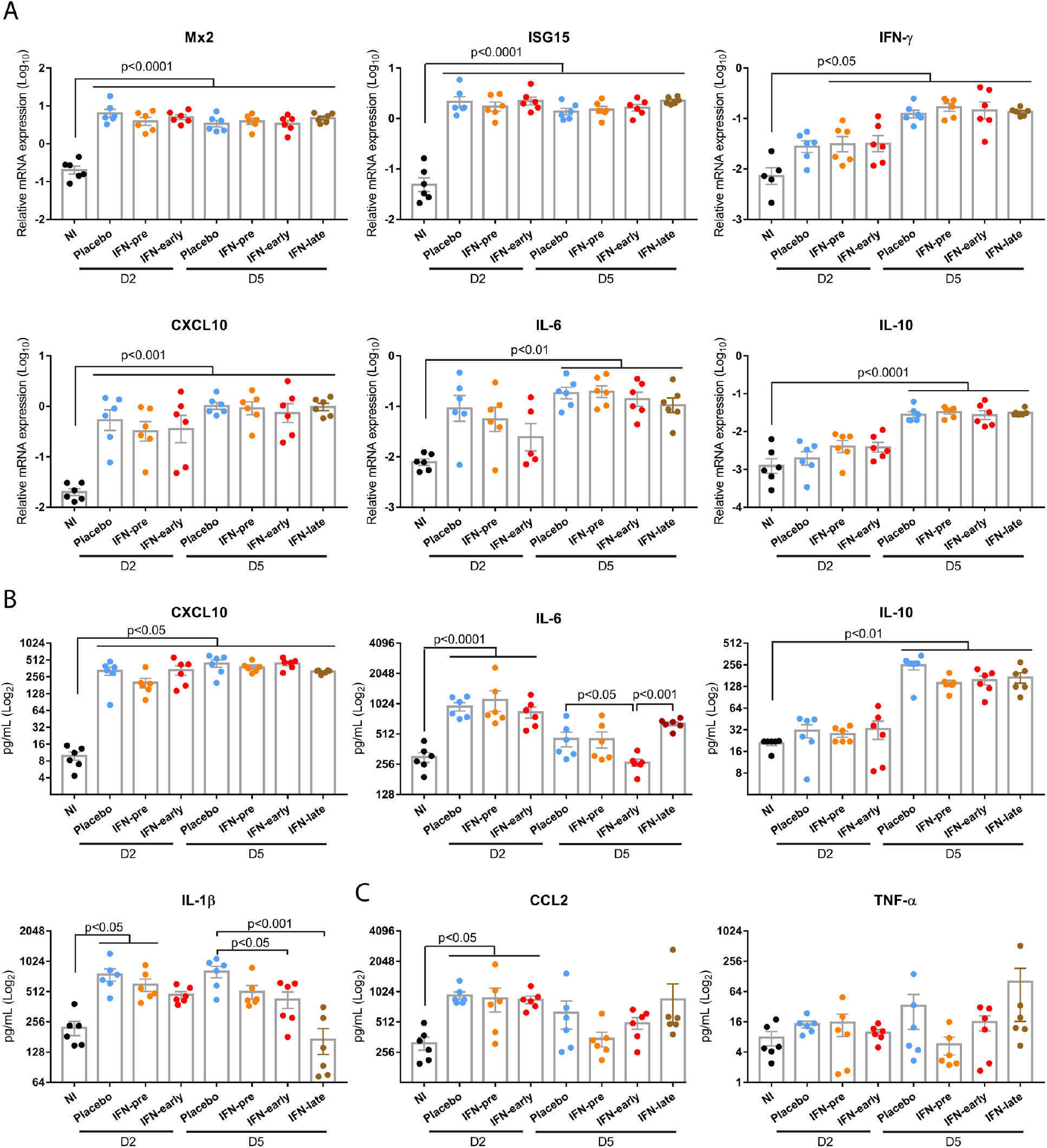
Impact of type I IFN-α treatment on the immune response to SARS-CoV-2. **(A)** Lung transcripts levels of Mx2, ISG15, IFN-γ, CXCL10, IL-6 and IL-10 relative to the housekeeping genes RPL18 and RPS6KB1 determined by RT-qPCR. **(B)** Lung protein levels for CXCL10, IL-6, IL-10 and IL-1β protein levels determined by ELISA or a multiplex assay. **(C)** Plasmatic protein levels for CCL2 and TNF-α determined by a multiplex assay. D2: day 2 post infection; D5: day 5 post infection; D15: day 15 post infection. Results are expressed as means ± SEM.

In this study, we assessed the *in vivo* prophylactic and therapeutic efficacy of type I IFN treatment against SARS-CoV-2 infection in the hamster model. Our study demonstrates that type I IFN treatment is beneficial when administered prophylactically or one day post-infection, confirming recent observations in hamsters (*20*). Similar findings were obtained when mice were treated prophylactically or at 12 hours post-infection with type III interferon or with the synthetic viral RNA analog poly(I:C) (*21, 22*). Altogether, these studies demonstrate that stimulation of the antiviral innate immune response before infection or at the very early phase of infection inhibits SARS-CoV-2 replication and pathogenesis, as expected given the high level of SARS-CoV-2 sensitivity to prophylactic type I and type III IFN treatment observed in cell culture (*2–4*). By contrasts, administration of type I IFN as soon as the animals exhibited the first clinical signs, corresponding to weight loss, three days post-infection, was not associated with any change in clinical signs compared to placebo treated hamsters. Therefore, this study does not support the use of intranasal type I IFN as a therapeutic in patients with COVID-19 symptoms.

In comparison to humans, virus replication and lung pathology progress much faster in hamsters, which have a peak of virus replication in the lungs at day 2-3 and a peak in clinical signs at day 6-7 (*17*). Treatment at day 3 post-infection thus corresponds to a “late” time point for treatment initiation in hamsters. We detected an upregulation of IL-6, CCL2 and TNF-α protein levels in the IFN-late group, compared to the IFN-pre and IFN-early groups. However, this did not result in enhanced pathology compared to the placebo group. Our results therefore indicate that type I IFN treatments at late time-points are unlikely to be associated with deleterious immunopathology exacerbation mechanisms, which were observed in SARS-CoV-1 and MERS virus infected mice and feared in SARS-CoV-2-infected humans (*13, 15, 16*). A contribution of type I IFNs to SARS-CoV-2 lung pathology has been suggested from work in IFNAR knockout mice transduced with adenovirus expressing human ACE2 and in STAT2 knockout hamsters (*21, 23, 24*). However, heightened viral loads were also observed in IFNAR knockout mice and STAT2 knockout hamsters, illustrating the fact that type I IFNs can have beneficial and deleterious effects most likely depending on the stage of SARS-CoV-2 infection (*21, 24*). SARS-CoV-2 expresses a broad array of type I IFN signaling antagonists that likely account for the low sensitivity observed in post-infection type I IFN treatments in cell culture (Fig. S8), and for the lack of beneficial effects observed in this study when type I IFN treatment was initiated at the onset of symptoms three days post-infection (*4, 25, 26*).

Even though SARS-CoV-2 expresses a broad array of type I IFN signaling antagonists, we detected a significant upregulation of ISGs expression in the respiratory tract of SARS-CoV-2 infected hamsters, similarly to what has been described in COVID-19 patients (*27*). Interestingly, ISGs expression in the respiratory tract was not further increased by IFN treatments, as previously observed in MERS-CoV infected mice treated with IFN-beta at 2 days post-infection (*16*). This result suggests that ISGs levels had reached their maximal in response to virus-induced endogenous type I and type III IFNs production and could not be further augmented following exogenous type I IFN administration.

Our study demonstrates that the timing of the type I IFN treatment is critical for its efficacy in a preclinical model of severe SARS-CoV-2 infection. Results from the SOLIDARITY clinical trial showed no benefit of subcutaneous interferon-β-1a injection, while a phase-two clinical trial provided evidence of some benefits of inhaled interferon-β-1a in COVID-19 patients (*12, 28*). The route of type I IFN administration was not the sole difference between these trials, as the patients treated in the SOLIDARITY trial were on average at a more severe stage of the disease. Our findings support the hypothesis that type I IFN treatments may only be beneficial in patients with mild symptoms at the early stages of the disease, while they are likely to provide no benefit in COVID-19 patients requiring hospitalization (*29*).

## Acknowledgments

We thank Meriadeg Ar Gouilh and Astrid Vabret (University Hospital of Caen, Normandie Université, Caen, France) for granting access to the SARS-CoV-2 UCN1 strain and Daniel Gonzalez-Dunia for proofreading the manuscript. We thank Vanessa Bastid, Mélanie Badré-Biarnais, Jean-Marc Boucher, Anouck Labadie, Carine Peytavin de Garam, Jonathan Rieder and Jean-Luc Schereffer for their investment in virological and serological analyses; Valère Brogat, Sébastien Kempff and Estelle Litaize for animal care and experimentation (Nancy laboratory for rabies and wildlife, ANSES, Lyssavirus Unit, Malzéville, France).

## Funding

this work was funded by a grant from the Agence Nationale de la Recherche (ANR-20-COV5-0004).

## Author contributions

R.V., G.F., P.B. and E.M.L. conceptualized and designed experiments. P.B., M.W., E.P.M., A.S., T.F., C.F.L., A.C., S.L., F.B., N.C., B.G., C.T., F.J.S., E.M.L. and R.V. performed experiments. R.V., P.B., F.J.S., G.F. and E.M.L analysed data. R.V., and P.B. wrote the paper. R.V. acquired funding.

## Competing interests

Authors declare no competing interests.

## Data and materials availability

All data is available in the main text or the supplementary materials.

## Supplementary Materials

Materials and Methods

Figures S1-S9

Tables S1

References (*30–38*)

## Notes

### Competing Interest Statement

The authors have declared no competing interest.

