## Supplementary materials for "Intranasal type I interferon treatment is beneficial only when administered before clinical signs onset in the SARS-CoV-2 hamster model"

**This PDF file includes:**

Materials and Methods
Figs. S1 to S8
Tables S1

### Materials and Methods

Animals and interferon-α treatment.

The animal experimentation protocols complied with the regulation 2010/63/CE of the European Parliament and of the council of 22 September 2010 on the protection of animals used for scientific purposes. These experiments were approved by the Anses/ENVA/UPEC ethic committee and the French Ministry of Research (Apafis n°24818-2020032710416319). 78 eight week-old female Syrian golden hamsters (*Mesocricetus auratus*, strain RjHan:AURA) were purchased from Janviers's breeding Center (Le Genest, St Isle, France) and housed in an animal-biosafety level 3 (A-BSL3), with *ad libidum* access to water and food. Animals weighing on average 97 grams at 1 day before infection (dbi) were randomly assigned to five groups: 12 non-treated non-infected animals (NI), 18 placebo-treated infected animals (Placebo), 18 infected animals interferon (IFN)-α-treated at 16 hours before infection (hbi), 1 day post-infection (dpi) and 3 dpi (IFN-pre), 18 infected animals IFN-treated at 1 and 3 dpi (IFN-early), and 12 infected animals IFN-treated at 3 and 5 dpi (IFN-late). At day 1, 3 and 5, all animals were anesthetized with isoflurane and treated by the intranasal route in each nostril either with 75µL PBS (Placebo) or with 75µl PBS containing $10^5$ UI of recombinant universal IFN-α (rIFN) (Hu-IFN-αA/D[Bg/II], pbl assay science, Piscataway, NJ). Animals from group IFN-pre were also anesthetized and IFN-treated 1 day prior to infection. In a preliminary experiment designed to test the efficacy of recombinant universal IFN-α in Syrian hamsters, eight week-old female Syrian golden hamsters (*Mesocricetus auratus*, strain RjHan:AURA), purchased from Janviers's breeding Center (Le Genest, St Isle, France) were treated intranasally with 75µl PBS containing $10^5$ UI of recombinant universal IFN-α (rIFN) (Hu-IFN-αA/D[Bg/II], pbl assay science, Piscataway, NJ) in each nostril. 12 hours post-treatment the animals were euthanized to harvest tissues for gene expression analyses.

Virus and experimental infection.

SARS-CoV-2 strain UCN1 was amplified as described previously and used at passage 2 (*30*). The viral stock was sequenced by Eurofins Genomics (Ebersberg, Germany) using the Illumina deep sequencing Eurofins Genomics Covid Pipeline v.0.1. Sequence analysis revealed that the virus had an intact spike cleavage site. Animals were anesthetized with isoflurane and intranasally inoculated with 20µL containing $5x10^3$ plaque forming units of UCN1 SARS-CoV-2 strain in each nostril. Non-infected animals received the equivalent amount of PBS. Animals were weighted daily from 1 dbi to 15 dpi. Oro-pharyngeal swabs were performed daily from 1 dpi to 6 dpi and at 8, 10 and 12 dpi. Six animals from groups Placebo, IFN-pre and IFN-early were anesthetized and euthanized by exsanguination at 2 dpi and then necropsied. Six animals from each group were also necropsied at 5 dpi. All remaining animals were necropsied at 15 dpi. For each necropsied animal, the following samples were collected: EDTA whole blood, lungs, spleen and nasal turbinates. Organs were either stored frozen at -80 °C in TRIzol reagent (Invitrogen, Carlsbad, CA) or Dulbecco's Modified Eagle medium (DMEM) containing penicillin and streptomycin, or stored in 10% neutral formalin.

Histology.

Samples from lung, upper trachea and larynx were fixed by immersion in 10% neutral-buffered formalin and processed routinely into paraffin wax. 4 µm sections were cut and stained with haematoxylin and eosin (H&E). In addition, samples were stained using the RNAscope in-situ hybridization (ISH) technique to identify the SARS-CoV-2 virus RNA, as previously described (31). Briefly, tissues were pre-treated with hydrogen peroxide for 10 minutes (room temperature), target retrieval for 15 mins (98-101°C) and protease plus for 30 mins (40°C) (Advanced Cell Diagnostics). A V-nCoV2019-S probe [Cat No. 848561, Advanced Cell Diagnostics] was incubated on the tissues for 2 hours at 40°C. Amplification of the signal was carried out following the RNAscope protocol using the RNAscope 2.5 HD Detection kit – Red (Advanced Cell Diagnostics). Tissue sections were also stained using immunohistochemistry (IHC) to detect Mx1 using a monoclonal antibody (Merck Sigma-Aldrich MABF938 clone M143/CL143) at 1:1,000. Tissues were dewaxed before heat-induced epitope retrieval was performed using Leica ER1 (pH 6.0) solution for 20 minutes on the Leica Bond RXm automatic stainer. Tissues were treated with hydrogen peroxide (5 mins) and a universal blocker (Superblock TBS blocking buffer, Thermo Scientific; 15 mins) before incubating the antibody for 15 mins. The Leica Bond Polymer Refine detection kit was used for visualisation and counterstaining. Tissue slides were scanned with a Hamamatsu Nanozoomer S360 scanner, visualized with NDP.view2 software and examined by a veterinary pathologist blind to the experimental conditions. A semi-quantitative analysis was used to evaluate the severity of lesion in the sections, as 0=none; 1=minimal; 2=mild; 3=moderate and 4=marked/severe. Digital image analysis Nikon-NIS-Ar software (version 4.30.01) was used to calculate the percentage of area with lesion and with positive staining in ISH and IHC slides.

Hematology.

For each necropsied animal, a complete blood count was performed within 15 minutes of sampling on a ProCyte Dx analyser (IDEXX laboratories, Westbrook, ME). Blood films were also performed, air-dried, stained with May-Grünwlad-Giemsa stain, fixed and coverslip-mounted. They were examined by a board-certified veterinary pathologist, blinded to the experimental conditions, to estimate the leukocyte differential count. The percentages of neutrophils, lymphocytes, monocytes, eosinophils and basophils were estimated from 100 cells. Samples with blood clots were excluded from the hematological analysis.

RNA extraction from tissue samples and cDNA synthesis.

For each organ, 30 mg portions of tissue were placed in tubes with beads (Precellys lysis kit; Stretton Scientific, Ltd., Stretton, United Kingdom) filled with 500 µL of TRIzol reagent (Invitrogen, Carlsbad, CA) and mixed for 5 s at 6,000 rpm three times in a bead beater (Precellys 24; Bertin Technologies, Montigny-le-Bretonneux, France). After TRIzol extraction, the aqueous phase was transferred to 96-well plate and processed according to the manufacturer's instructions (NucleoMag RNA; Macherey-Nagel GmbH & Co, Germany) using a KingFisher automated platform (Thermo Fisher Scientific, Inc., Ontario, Canada). cDNA was synthesized by reverse transcription of 500 ng of total RNA using both oligo(dT)18 (0.25 µg) and random hexamer (0.1 µg) and a

RevertAid first-strand cDNA synthesis kit (Invitrogen, Thermo Fisher Scientific) according to the manufacturer's instructions.

Quantitative PCR from tissue samples.

Quantitative PCR for the analysis of host genes expression was performed in 96-well plates in a final volume of 10 µL on a LightCycler 96 (Roche, Mannheim, Germany). Mixes were prepared according to the manufacturer's instructions (QuantiFast SYBR green PCR; Qiagen) with 1 µL of 1:2 diluted cDNA and a final 1 µM concentration of each primer (Table S1). Relative quantification was carried out by using the $2^{-\Delta CT}$ method, and the geometric means of two housekeeping genes (RPL18 and RPS6KB1).

Quantitative PCR from oropharyngeal swabs.

Viral RNA was extracted from 160 µL of oro-pharyngeal swabs stored in 500µL of DMEM with antibiotics. RNA extraction was performed by using Qiagen Viral RNA mini kit according to the manufacturer's instructions (Qiagen, Les Ulis, France), with the addition of 15µL Triton X-100 (MP Biomedicals, Illkirch, France) to 560 µL of AVL Lysis buffer for each sample. TaqMan RT-qPCR was performed according to the manufacturer's instruction (QuantiTect Probe RT-PCR; Qiagen) using primers targeting the envelope protein gene (E gene) and a previously described protocol (*32*). Absolute quantification was performed using a standard curve based on six 10-fold dilutions of a quantitative Synthetic RNA from SARS-CoV-2 (BEI Resources : Catalog No. NR-52358).

IL-6 ELISA from lung samples.

150 mg portions of lung tissue were placed in tubes with beads (Precellys lysis kit; Stretton Scientific, Ltd., Stretton, United Kingdom) filled with 1 mL of Dulbecco's Modified Eagle medium (DMEM) and mixed for 5 s at 6,000 rpm three times in a bead beater (Precellys 24; Bertin Technologies, Montigny-le-Bretonneux, France). Supernatant was collected after a brief centrifugation and a commercial hamster IL-6 double antibody enzyme-linked immunosorbent assay kit was performed according to the manufacturer's instructions (ELISA Genie, Dublin, Ireland).

Cytokine/chemokine quantitation using a multiplex assay

150 mg portions of lung tissue were placed in tubes with beads (Precellys lysis kit; Stretton Scientific, Ltd., Stretton, United Kingdom) filled with 1 mL of Dulbecco's Modified Eagle medium (DMEM) containing Triton X100 (1% v/v to inactivate the virus) and a protease inhibitor cocktail (Sigma). Tubes were mixed for 5 s at 6,000 rpm three times in a bead beater (Precellys 24; Bertin Technologies, Montigny-le-Bretonneux, France). Plasma samples also contained Triton X100 (1% v/v to inactivate the virus) and a protease inhibitor cocktail (Sigma). Cytokines and chemokines concentrations were determined using a custom developed Syrian Hamster cytokines MilliPlex xMAP assay (Merck Millipore) following the manufacturer instructions. Data were recorded on a MagPix instrument using Xponent software (Luminex). Results are expressed as concentration in pg/mL. Among the cytokines and chemokines included in the multiplex assay, we detected a robust signal for CXCL10, IL-10, IL-1β in lung and plasma samples.

However, CCL2 and TNF-α were only detected in plasma samples, most likely due to higher signal to noise ratio in plasma samples than in lung samples.

*In vitro* IFN-α treatment

Vero-E6 cells were grown in DMEM containing 1% antibiotics (penicillin/streptomycin) and 10% fetal bovine serum. Cells were treated with 1000 UI/mL recombinant universal IFN-α (rIFN) (Hu-IFN-αA/D[Bg/II], pbl assay science, Piscataway, NJ) for 18 hours prior to infection and were subsequently infected with SARS-CoV-2 at a multiplicity of infection (MOI) of $10^{-3}$. Infections were carried out in DMEM containing 1% antibiotics (penicillin/streptomycin) and 2% fetal bovine serum. Recombinant universal IFN-α (Hu-IFN-αA/D[Bg/II], pbl assay science, Piscataway, NJ) was added at a final concentration of 1000 UI/mL to the medium 6 hours post-infection for cells treated subsequently to the infection. Culture supernatants were harvested 24 hours post-infection and viral titers were determined by the $TCID_{50}$ method on Vero-E6 cells and calculated by the Spearman & Kärber algorithm.

SARS-CoV-2 titrations from lung samples.

Titrations were performed on 90% confluent Vero-E6 cells in 96-well plates. Viral titers were calculated using the Spearman-Kärber method.

Statistical analyses

All the data, except weight loss, were statistically analyzed using one-way analysis of variance (ANOVA) followed by Tukey's post hoc test (GraphPad Prism Software, USA) with the p values indicated corresponding to the results of Tukey's post hoc tests. Weight loss was statistically analyzed using two-way ANOVA with Geisser-Greenhouse correction followed by Dunnet's multiple comparisons test (GraphPad Prism Software, USA) with the p values indicated for the weight loss corresponding to the results of Dunnet's post hoc tests.

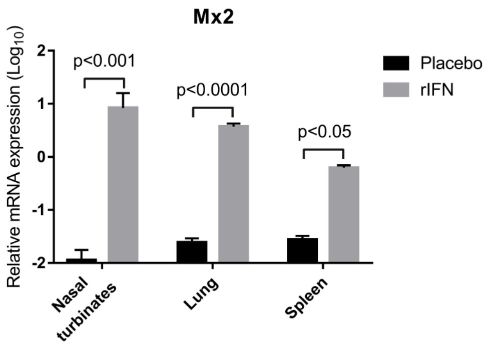


**Fig. S1. Impact of IFN-α treatment on Mx2 transcript levels in non-infected hamsters.**

Syrian hamsters were treated intranasally either with placebo or with $2x10^5$ UI recombinant universal IFN-α (rIFN)**.** Tissues were harvested 12 hours post-treatment. Transcripts levels of Mx2 relative to the housekeeping genes RPL18 and RPS6KB1 were determined by RT-qPCR. Results are expressed as means ± SEM.

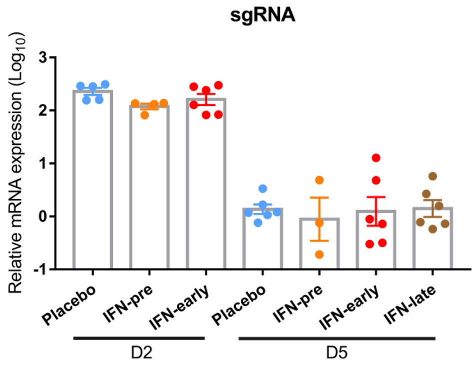


**Fig. S2. Subgenomic viral RNA in nasal turbinates.**

Nasal turbinates were harvested at day 2 post-infection (D2) or day 5 post-infection (D5). Viral sgRNA levels relative to the housekeeping genes RPL18 and RPS6KB1 were determined by RT-qPCR. Results are expressed as means ± SEM.

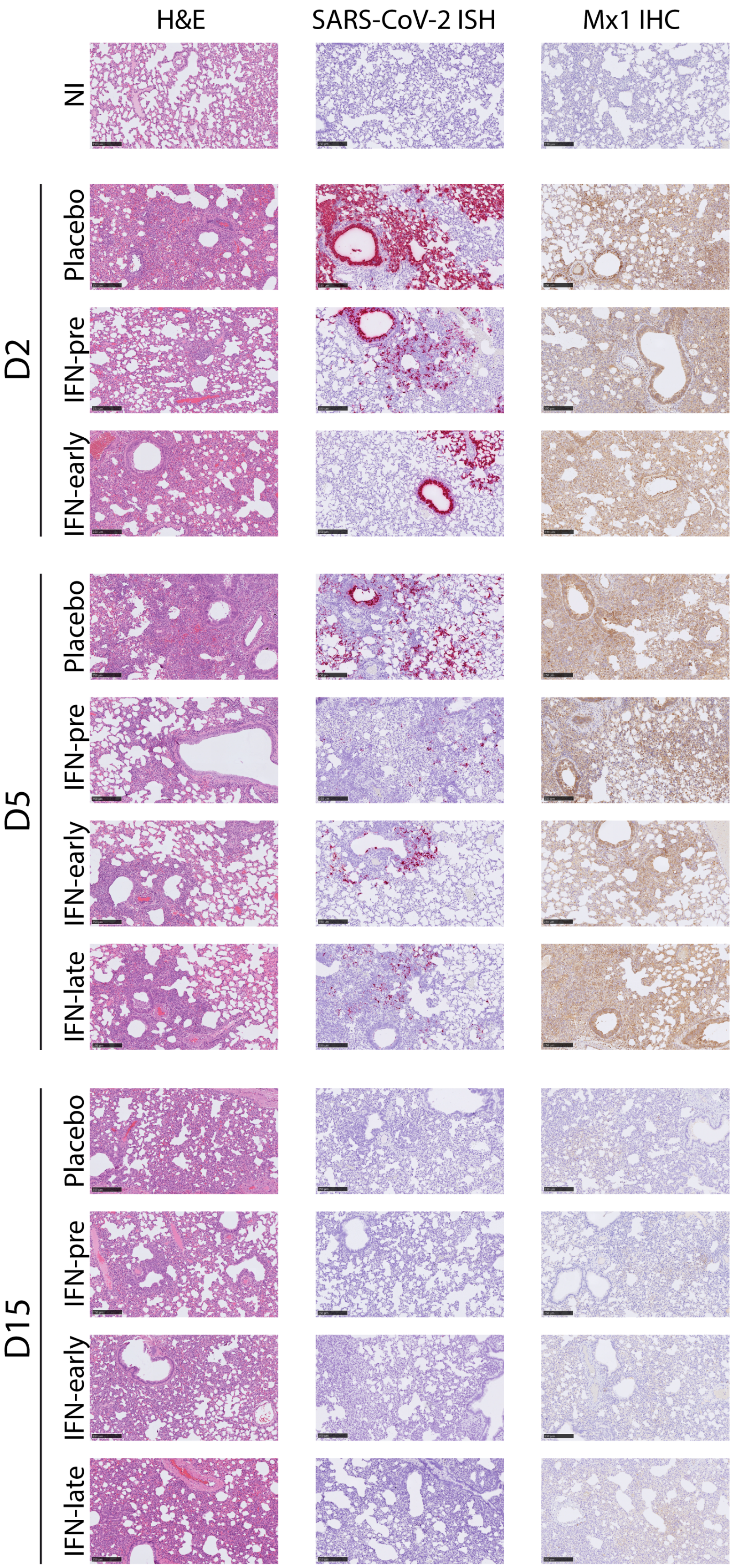
H&E
SARS-CoV-2 ISH
Mx1 IHC
NI
D2
Placebo
IFN-pre
IFN-early
D5
Placebo
IFN-pre
IFN-early
IFN-late
D15
Placebo
IFN-pre
IFN-early
IFN-late

**Fig. S3. Histological analysis of the impact of IFN-α treatments.**

Representative pictures were selected to display the pathology from haematoxylin and eosin (H&E) stained lung section, viral RNA in lung sections stained with RNAScope *in situ* hybridization (ISH) and Mx1 protein detected by immunohistochemistry (IHC). D2: day 2 post infection; D5: day 5 post infection; D15: day 15 post infection.

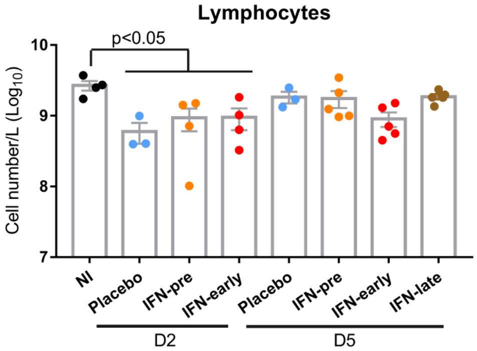


**Fig. S4. Type I IFN-α treatment does not prevent lymphocytopenia.**

A complete blood count analysis was performed as described in the methods section. D2: day 2 post infection; D5: day 5 post infection. Results are expressed as means ± SEM.

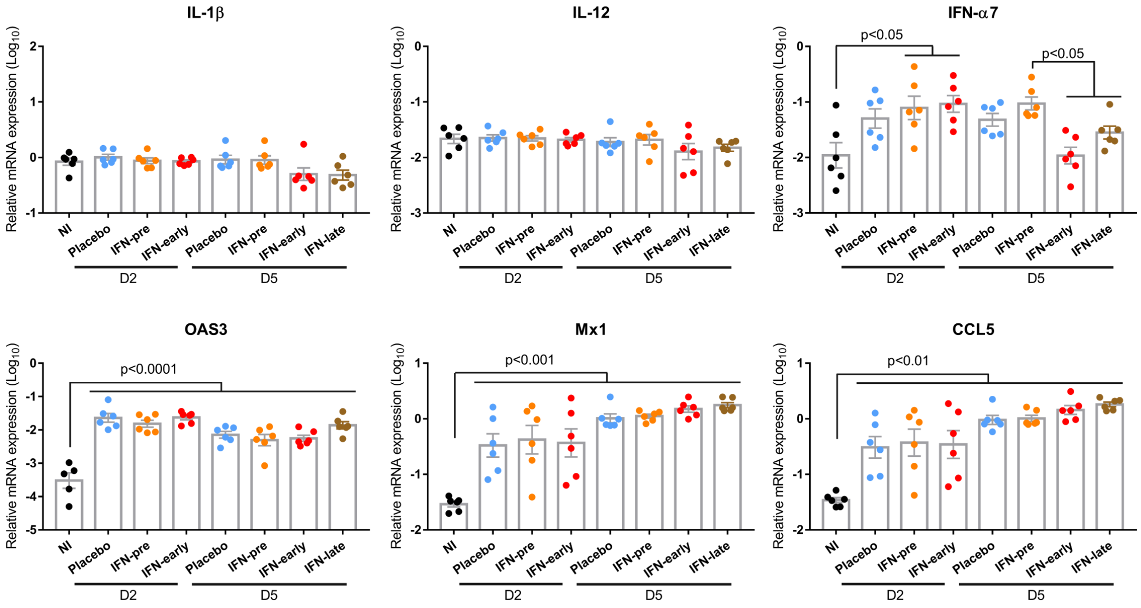


**Fig. S5 . Impact of type I IFN-α treatment on the immune response to SARS-CoV-2. in the lung.**

Lung transcripts levels of IL-1β, IL-12, IFN-α7, OAS3, Mx1, CCL5 relative to the housekeeping genes RPL18 and RPS6KB1 determined by RT-qPCR. D2: day 2 post infection; D5: day 5 post infection; D15: day 15 post infection. Results are expressed as means ± SEM.

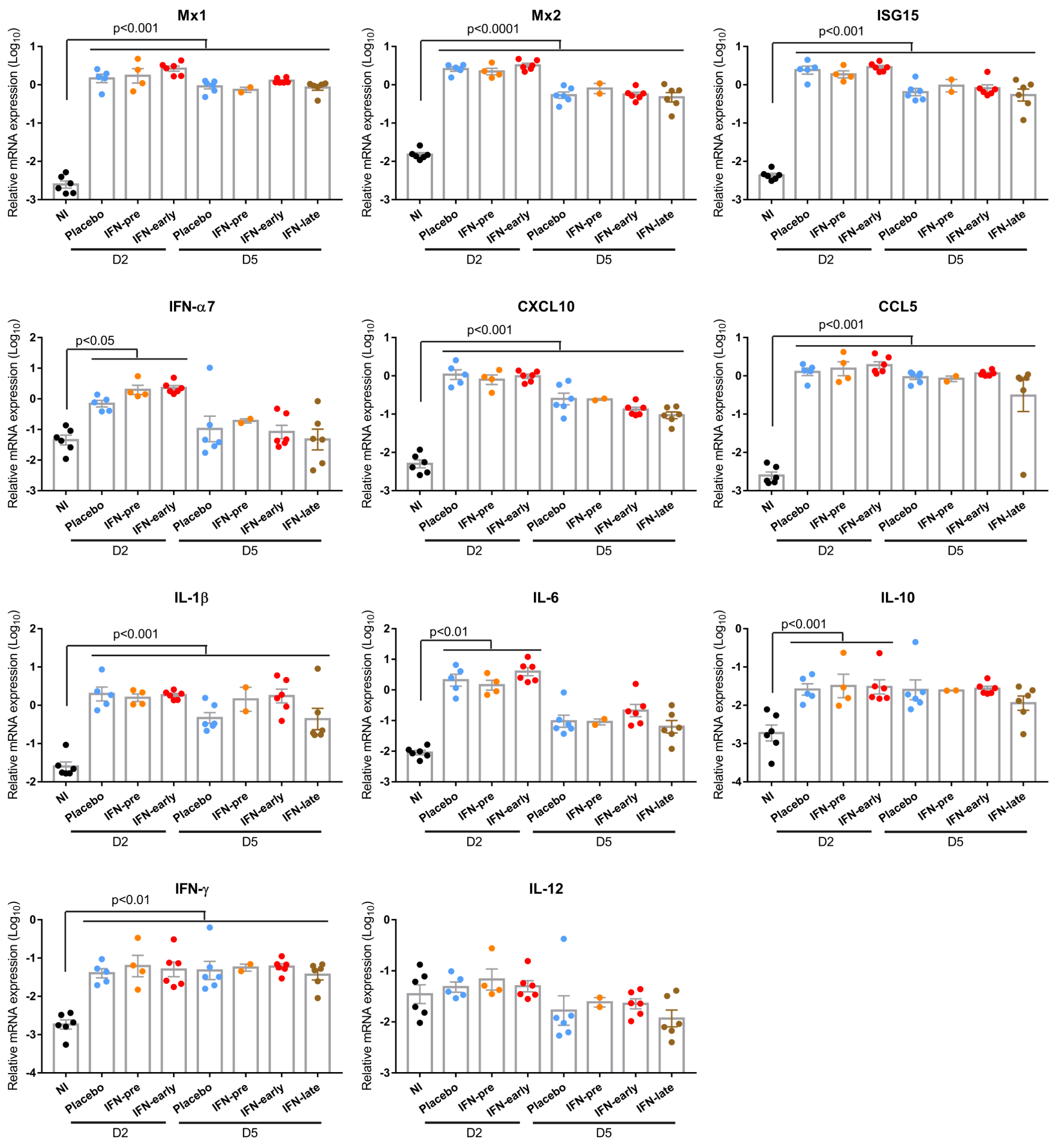


**Fig. S6. Impact of IFN-α treatment on the immune response to SARS-CoV-2. in the nasal turbinates.**

Nasal turbinates transcripts levels of Mx1, Mx2, ISG15, IFN-α7, CXCL10, CCL5, IL-1β, IL-6, IL-10, IFN-γ and IL-12 relative to the housekeeping genes RPL18 and RPS6KB1 determined by RT-qPCR. D2: day 2 post infection; D5: day 5 post infection; D15: day 15 post infection. Results are expressed as means ± SEM.

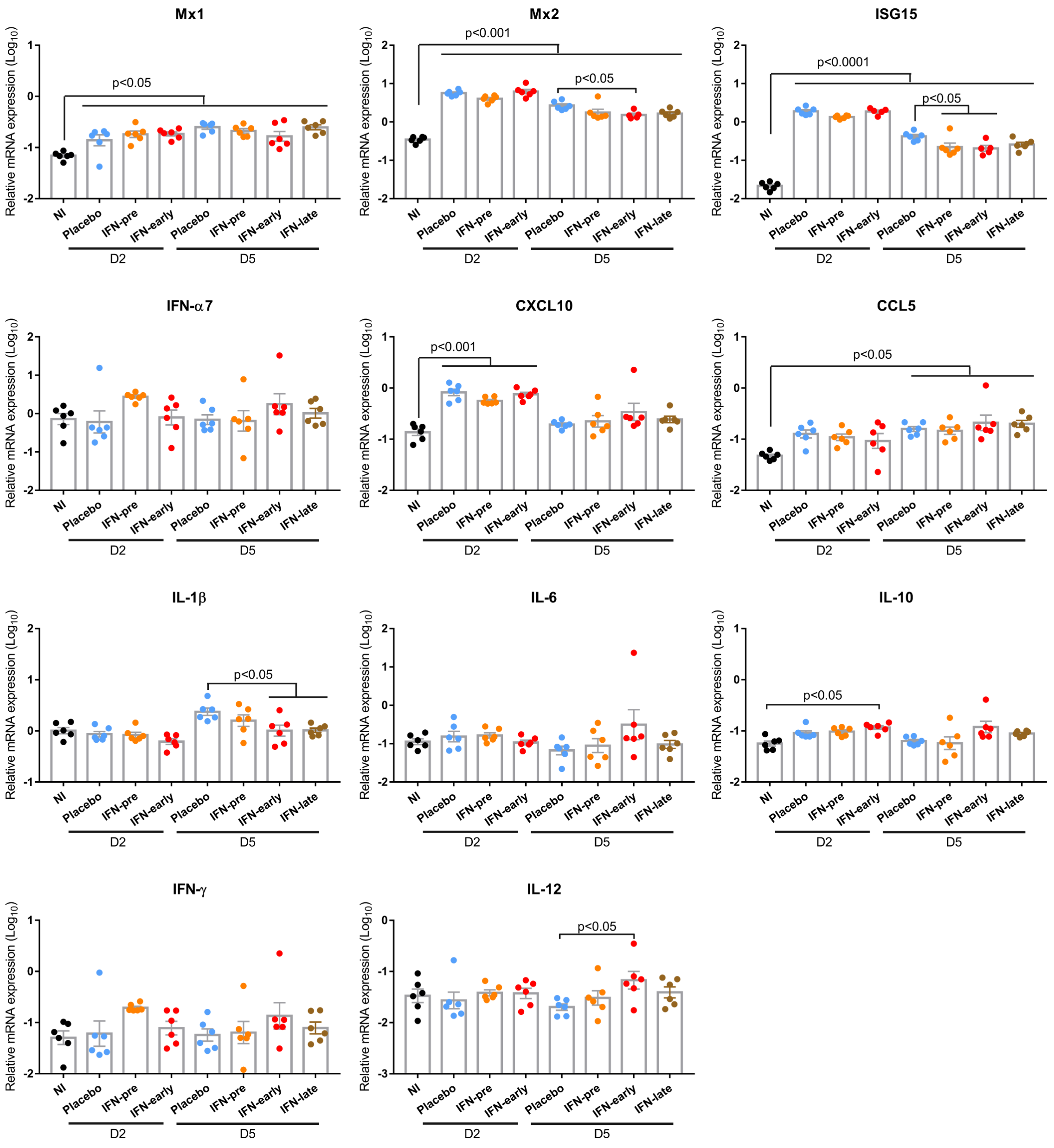


**Fig. S7. Impact of IFN-α treatment on the immune response to SARS-CoV-2. in the spleen.**

Spleen transcripts levels of Mx1, Mx2, ISG15, IFN-α7, CXCL10, CCL5, IL-1β, IL-6, IL-10, IFN-γ and IL-12 relative to the housekeeping genes RPL18 and RPS6KB1 determined by RT-qPCR. D2: day 2 post infection; D5: day 5 post infection; D15: day 15 post infection. Results are expressed as means ± SEM.

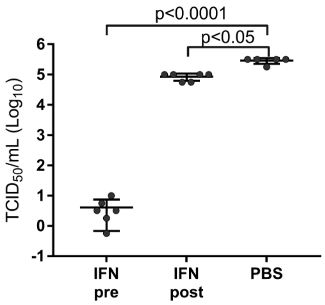


**Fig. S9. Effect of IFN-α treatment on SARS-CoV-2 replication in cell culture.**

Vero-E6 cells were treated either with placebo or with $10^3$ UI/mL recombinant universal IFN-α 18 hours prior to infection (IFN-pre) or 6 hours post infection (IFN-post). Equivalent volume of PBS was used as a negative control. Viral titers were determined by $TCID_{50}$ from supernatants collected 24 hours post infection. Each dot represents a technical replicate of a representative experiment performed twice. Results are expressed as means ± SEM.

**Table S1. List of primers**

| Gene name | Primers sequences (5' to 3') | NCBI accession number or Reference |
|---|---|---|
| CCL5 | ACTGCCTCGTGTTCACATCA | XM_005076936.3 |
| | CCTTCGGGTGACAAAAACGA | |
| CXCL10 | GCCATTCATCCACAGTTGACA | (33) |
| | CATGGTGCTGACAGTGGAGTCT | |
| IFN-α7 | CTGGTGGCTGTGAGGAAATA | (34) |
| | AGCAAGTTGGCTGAGGAAGA | |
| IFN-γ | GGCCATCCAGAGGAGCATAG | (35) |
| | TTTCTCCATGCTGCTGTTGAA | |
| IL-1β | GGCTGATGCTCCCATTCG | (36) |
| | CACGAGGCATTTCTGTTGTTCA | |
| IL-6 | CCTGAAAGCACTTGAAGAATTCC | (33) |
| | GGTATGCTAAGGCACAGCACACT | |
| IL-10 | GTTGCCAAACCTTATCAGAAATGA | (36) |
| | TTCTGGCCCGTGGTTCTCT | |
| IL-12 | GGCCTTCCCTGGCAGAA | (33) |
| | ATGCTGAAAGCCTGCAGTAGAAT | |
| ISG15 | AAAGCCTACAGCCATGACCT | XM_013119951.2 |
| | TTAGTCAGGGGCACCAGGAA | |
| Mx1 | GCGCTTCCAGACTCTTCTGA | XM_021229467.1 |
| | CCTAAGATACATGCGATGGCG | |
| Mx2 | CCAGTAATGTGGACATTGCC | (35) |
| | CATCAACGACCTTGTCTTCAGTA | |
| OAS3 | AGGTGCTTAAGGTGGTTAAGGG | (37) |
| | TGCTCAGAGAAGTGCTGGAAG | |
| RPL18 | GTTTATGAGTCGCACTAACCG | (35) |
| | TGTTCTCTCGGCCAGGAA | |
| RPS6KB1 | TCAGACCGGTGGAAAACTCTAC | (37) |
| | TGATGCAAATGCCCCAAAGC- | |
| SARS-CoV-2 TRS-L | CTCTTGTAGATCTGTTCTCTAAACGAAC | (38) |
| SARS-CoV-2 TRS-N | GGTCCACCAAACGTAATGCG | |
| TNF-α | GGAGTGGCTGAGCCATCGT | (33) |
| | AGCTGGTTGTCTTTGAGAGACATG | |
